# Platform for Closed Loop Neuromodulation Based on Dual Mode Biosignals

**DOI:** 10.1101/163329

**Authors:** Khalid B. Mirza, Nishanth Kulasekeram, Simon Cork, Stephen Bloom, Konstantin Nikolic, Christofer Toumazou

## Abstract

Closed loop neuromodulation, where the stimulation is controlled autonomously based on physiological events, has been more effective than open loop techniques. In the few existing closed loop implementations which have a feedback, indirect non-neurophysiological biomarkers have been typically used (e.g. heart rate, stomach distension). Although these biomarkers enable automatic initiation of neural stimulation, they do not enable intelligent control of stimulation dosage. In this paper, we present a novel closed loop neuromodulation platform based on a dual signal mode that is detecting electrical and chemical signatures of neural activity. We demonstrated it on a case of vagus nerve stimulation (VNS). Vagal chemical (pH) signal is detected and used for initiatisng VNS and vagal compound action potential (CAP) signals are used to determine the stimulation dosage and pattern. Although we used the paradigm of appetite control and neurometabolic therapies, the platform developed here can be utilised for prototyping closed loop neuromodulation systems before adapting the final System-on-Chip (SoC) design.

## I. Introduction

Neuromodulation is a fast growing treatment paradigm for a number of diseases and conditions such as epilepsy, depression, obesity, inflammation, etc [1]. Apart from implementing it for drug resistant cases, the primary reason behind this is the ability for neuromodulatory therapies to enable a more precise intervention by targeting a certain nerve or brain area than drug therapies. Furthermore, the dosage can be tuned for each individual case [2].

Neuromodulation therapies can be divided into two therapeutic paradigms: *open loop* and *closed loop*. The stimulation dose control aspect of neuromodulation implants consist of answering two questions: *when* to stimulate and *how much*. Open loop neuromodulation therapies involve crude manual tuning of stimulation time and dosage by healthcare professionals based on factors such as patient vitals, discomfort or pain, etc [3]. *Closed loop* involves autonomous decisions based on physiological events, hence it tends to be more patientspecific. Closed loop neuromodulation therapies have been demonstrated to be more efficient than open loop [4], [5].

In this paper, we introduce a novel platform to implement an adaptive stimulation based on dual mode signalling i.e chemical and electrical, see Fig. 1. Our immediate aim is to use it to develop a closed loop VNS implant for apettite control, but it can be used as generic platform for developing adaptable neurostimulation systems. This system is able to address the two primary concerns of a closed loop implant by initiating stimulation based on the presence of specific chemical signature in the neural response to a specific physiological condition and controlling the stimulation dose based on CAPs elicited say during an interrogative low frequency stimulation

**Fig. 1.**
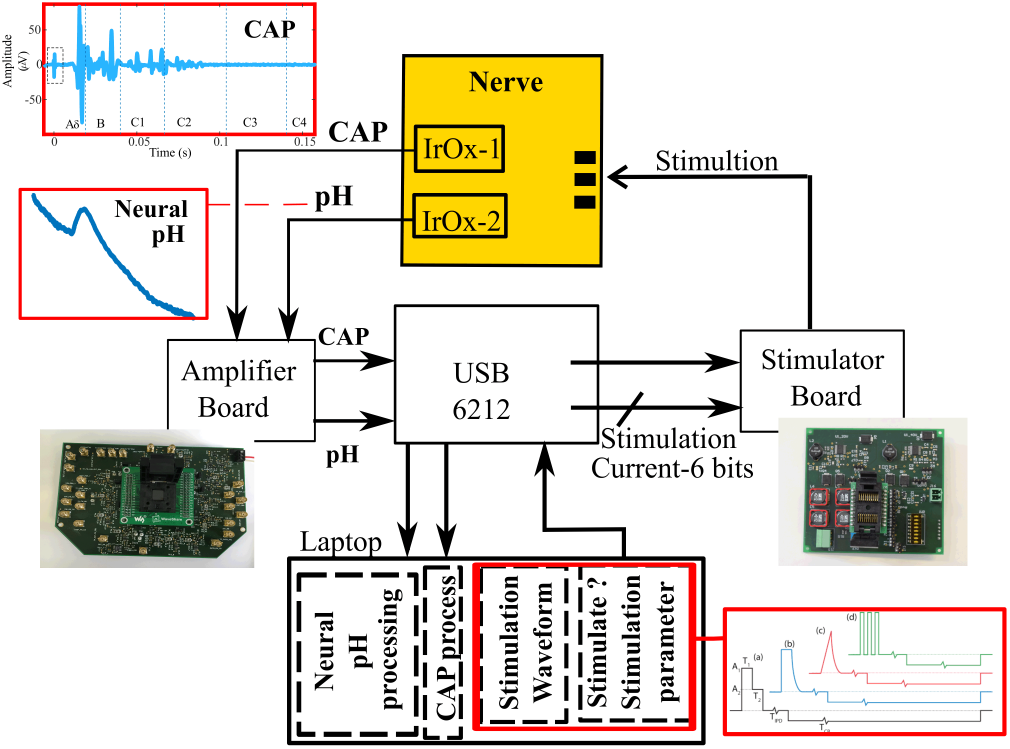
Platform for dual mode biosignal closed loop neuromodulation.

The platform was initially created for the development of a closed-loop stimulation of the gastric branch of the vagus nerve in order to regulate appetite [6]. We were able to identify chemical pH signatures specific to the vagus response to cholecystokinin (CCK), a gut hormone released during meal intake, during *in vivo* experiments [6]. CCK is responsible for reducing appetite and it has been previously demonstrated that VNS introduced in correlation to meal intake leads to greater effective weight loss [3]. We utilise the pH signal (chemical sensor) which has a higher amplitude, hence easy to detect compared to neural mass activity, to initiate VNS in correlation to meal intake. However, apart from knowing when to stimulate we needed to adjust the stimulation to a certain level which always varies from case to case due to electrodes proximity to the nerve, contact impedance etc. We utilise CAPs elicited during an interrogative VNS protocol to characterise the vagus nerve for individual subjects and determine the precise stimulation parameters necessary.

## II. System Architecture

The prototyping platform consists of all elements needed for a complete closed loop neuromodulation system: electrodes, front-end analogue amplifiers, signal processing algorithms and stimulator, Fig. 1. The electrodes are IrOx microspikes interfaced to a multichannel, multifunctional neural amplifier with on-chip 8-bit ADC. This ADC is interfaced to an NI USB 6212 Data Acquisition Device. The acquired neural signals are processed on the computer in this platform, with the future goal of enabling efficient System-On-Chip implementation. Finally, the PC is connected to a stimulator.

### A. Sensors: IrOx Electrodes

Choice of electrodes is a crucial part of the platform development. The instrumentation specification and design very much depend on it. Initially we have targeted to measure the pH, and currently we are examining potassium electrodes as well [7]. For pH we decided to use IrOx wires [8]. The mechanism behind ability of the IrOx to sense pH changes lies in the existence of redox chemical reaction between two IrOx types, namely Ir(III)Ox and Ir(IV)Ox:

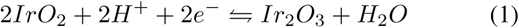

A potentiometric sensor typically measures potential difference between the sensing electrode and a reference electrode such as silver-silver chloride electrode (Ag/AgCl), i.e. the Open Circuit Potential (OCP). The measured OCP is modelled by the Nernst equation and is dependant on ionic concentration. The sensitivity of potentiometric IrOx sensors range from 60 − 90 mV/pH. For the typical pH variation observed in biomedical applications of 0.02 − 1 pH units and assuming minimum sensitivity, the signal levels will range from 1.2 mV to 60 mV. The response time for IrOx sensors i.e. time required to reach 90% of the equilibrium value, for the relevant ranges, is 6 − 12 s for a change of 1 pH unit [9]. Hence the pH signal is a low frequency signal with bandwidth less than 0.1 Hz.

An outstanding issue with IrOx sensors is presence of drift in the OCP. The origin of the drift could be due to hysteresis in the IrOx sensor or change in ambient conditions, such as temperature, which affect pH [9].

IrOx microneedles have small sensing area and in order to preserve this sensing layer, it is essential to ensure no leakage current flow through the electrode. Otherwise, this will lead to reduction of IrOx layer and hence loss of sensitivity. Therefore, it is crucial to use front-end amplifiers that have extremely low input bias current or gate leakage current in order to prevent sensor degradation and erroneous readings.

### B. Front-end Amplifier

The chip micrograph in Fig. 2A) shows the make up of the two varieties of front end amplifiers. The first is a modified switched bias amplifier (MSB), which was designed to exhibit a low input referred noise with a closed loop gain of 60 dB and it operates over a frequency range 200 Hz to 5 kHz. The MSB amplifier offers significant noise and area trade-off, all details are available in [10]. The second variety is a chemical amplifier based on the MSB amplifier with the closed loop gain clipped to a maximum of 20 dB and a maximum frequency range of 10 Hz. Each of these amplifiers have been arrayed to create a multichannel amplifier consisting of three MSB based amplifiers which are used to pickup CAP signals and the remaining three are used to pickup chemical based signals. The amplifiers were designed and fabricated in the standard 0.35 *μ*m CMOS AMS technology. The PCB, Fig 2B) was designed primarily to test the amplifiers mentioned above and interface with the rest of the platform. Due to the low power consumption it is capable of operating off a battery.

**Fig. 2.**
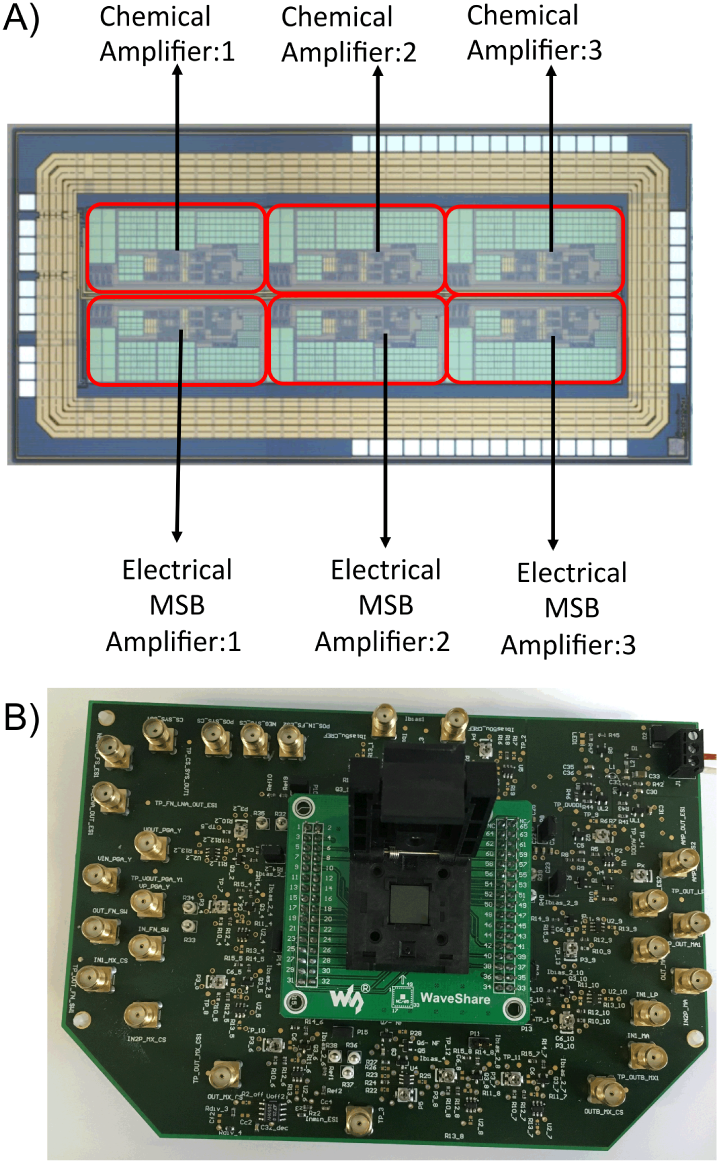
Multi-channel Recording setup: A) chip microphotograph, B) evaluation PCB.

### C. Neural Stimulator

The heart of the bi-phasic stimulator is a 6 bit current DAC, Fig. 3A). The DAC has a Least Significant Bit (LSB) of 50 *μ*A, and a maximum current output of 3.2 mA. To maintain charge balance a current leakage estimator is included as part of the circuitry. If there is a loss of charge when driving a current pulse from the anode through the nerve tissue to the cathode, an additional current pulse for the difference in charge, is sent to the driving terminals. The design of the bi-phasic stimulator was carried out using commercially available High Voltage AMS H35 technology which included a low voltage 0.35 *μ*m extension. A high voltage technology is needed to drive a 3.2 mA current into a tissue with a nominal impedance of 10 kΩ, resulting in a minimum voltage of 32 V, needing a power supply of approximately 35 V to drive the output stage of the bi-phasic stimulator.

**Fig. 3.**
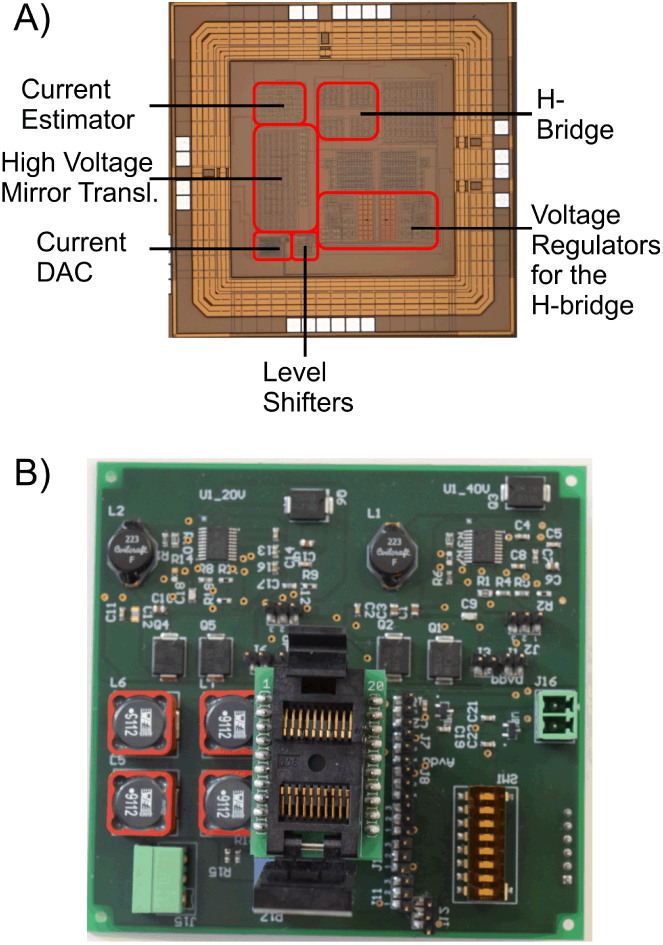
Neural Stimulatior: A) chip microphotograph, B) evaluation PCB.

As the high voltage transistors were relatively large compared to the low voltage transistors a decision was made to design the 6 bit current DAC using the low voltage extension.

The H-bridge was used to drive the anode and cathode terminals, and form the basis of the bi-phasic operation. High Voltage cascoded mirrors were designed to interface between the low voltage 6 bit current DAC and the H-bridge. Careful design consideration was given to the H-bridge high voltage transistors to ensure junction voltages do not exceed and cause fatigue of the oxide and eventual break down. To avoid such situations a high voltage regulator was designed to ensure junction voltages within the H-bridge doesn’t exceed specified typical voltage, as indicated in the design manual for the various transistor flavours.

The stimulator PCB, shown in Fig. 3B) was designed to interrogate the integrated stimulator IC and demonstrate that it is capable of producing 6 bit current output depending on the input digital code, with an LSB of 50 *μ*A. As the final solution is expected to be battery powered the PCB has a boost converter to up convert 3.3 V to 40 V, which is then dropped to the 35 V needed for driving the output stage of the stimulator integrated circuit.

### D. Closed Loop Implementation

The closed loop implementation consists of three different steps: *Nerve classification*, *Stimulation strength determination*, *Physiological Trigger Detection*, an overview shown in Fig. 4.

**Fig. 4.**
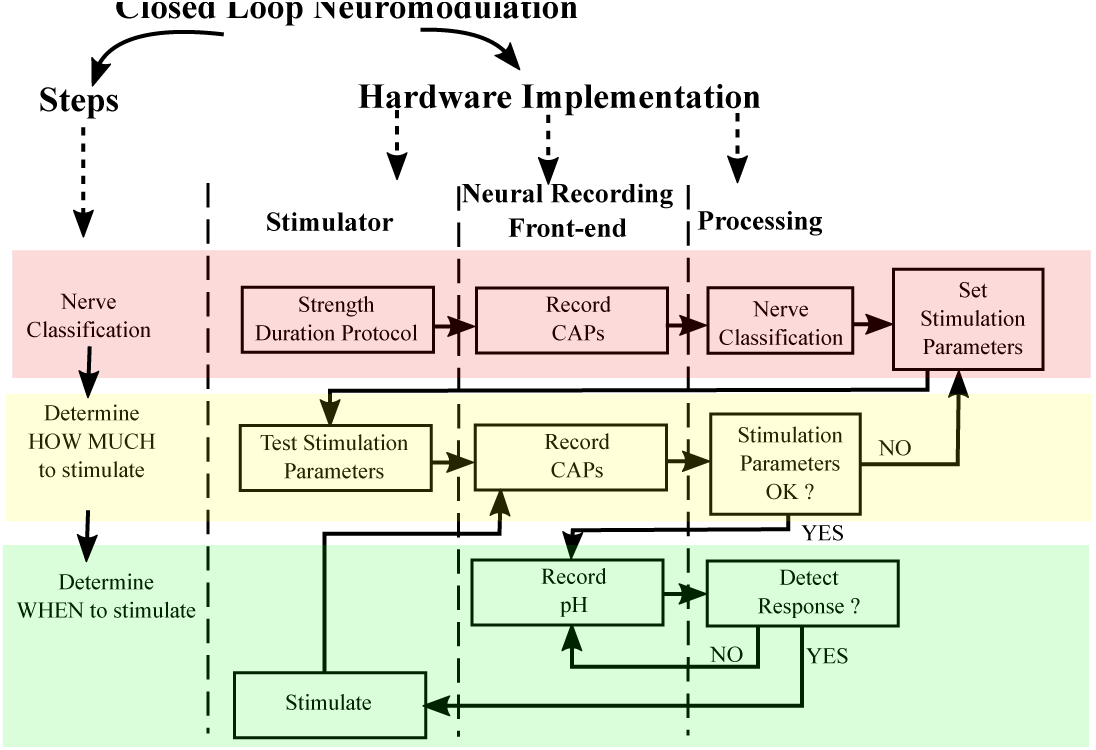
Overview of closed loop neuromodulation system.

*Nerve classification* is performed to determine the excitabilty of the nerve, including the types of fibres present and the stimulation thresholds for different fibre types. For this purpose we use the Strength–Duration protocol [11]. The CAP waveform analysis consists of partitioning the CAP waveform to separate the contirbution of different fibre types based on fibre conduction velocity and distance between stimulation and recording electrode [2]. A strength duration curve is constructed for each fibre type based on the Lapicque equation [2]. *Stimulation strength determination* is performed by setting the stimulation parameters i.e stimulation current, pulsewidth, and stimulation waveform based on the neuromodulatory application. A test stimulus is performed to verify the set stimulation parameters.

Once the stimulation parameters are verified, the next step is to set up the platform to detect the physiological trigger for initiating stimulation. In the application described here the physiological trigger is the pH change due to vagus nereve response induced by release of CCK. The incoming data which is sampled at 20 kHz is downsampled using a decimation filter to a 50 samples per second. The decimation filter is composed of a CIC filter and a compensation low pass filter.

*1) pH Electrode Drift Removal:* The chemical signal has an overall linear drift due to the drift in open circuit potential of the IrOx electrode as shown in Fig. 5. This drift is cancelled using linear interpolation over a fixed time window.

**Fig. 5.**
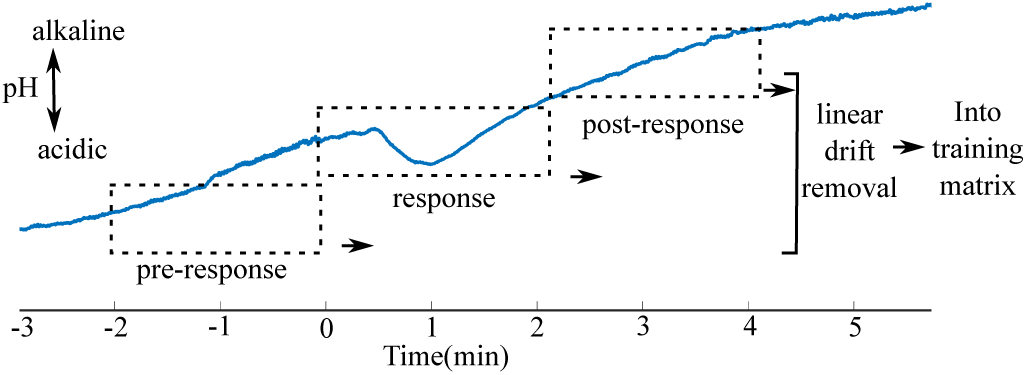
The pH change measured *vivo* in a rat model inside the gastric branch of the vagus nerve, before injection of CCK (*pre*), during the period when CCK is active (*response*) and after (*post-response*).

*2) pH Detection Algorithm Training:* In order to detect the CCK specific pH response, the temporal profile of the pH waveform is considered over a period of 2 minutess and compared with the CCK induced temporal pH profile. In general, the CCK specific pH response exhibits a negative slope or a downward trend for 1-1.5 minutes followed by a reverse trend for the same length of time, see Fig. 5. However, under *in vivo* conditions, the pH waveform is affected by several interefering processes. Hence, in this algorithm a multivariate approach is adopted by using Principal Component Analysis (PCA). This is similar to the application of PCA on cyclic voltammetry curves described in details in [12], [13].

PCA is performed by first mean centering the data and constructing a training matrix using *in vivo* experimental data, in which CCK was injected intravenously and the change due to CCK was recorded, consistently over a number of trials in different animals [6]. If necessary, this training can also be performed *chronically* on-chip in an implant or using the platform described in this paper during *in vivo* experiments.

The training matrix consists of *pre* (2 min), *response* (2 min) and *post* (2 min) CCK injection as shown in Fig. 5. The principal components (PC) for *pre*, *post* and *response* (*P*_train_) are extracted. A scree plot is used to determine the number of PCs to retain, which capture at least 95% of the variance in data. These will be used for real time detection of pH response. The training matrix is cross-validated with a set of exisiting *in vivo* experimental data not included in the training matrix. The extracted PCs are stored for real time pH response detection.

*3) Real Time pH Response Detection:* The incoming real time pH data is filtered and a projection matrix of the data in the principal component subspace is calculated:

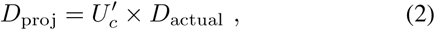

where *D*_actual_ is the real time experimental data and U_*c*_ is the Principal Component matrix extracted from the training matrix. The specific response to a physiological stimulus is detected by calculating *residual values E*, defined as the difference between the actual incoming data and back-transformed projected data set:

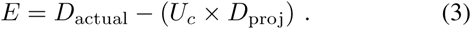

From residuals a factor *Q* is calculated [12]:

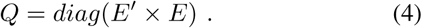

In other words *Q* is the sum of the squares of the residual value in each sample data. Now the value of the *Q* is used to determine the presence of a specific neural response by establishing a threshold value for *Q*. In our case, if the Q-value is less than 0.01, it is treated as a neural response.

In our case three minutes of real time data is stored at any given time. In order to capture the CCK response effectively, two projection matrices (temporal size 2 min) are calculated on 2 minutes of data with an overlap of 1 minute. Before calculating the projection matrices, electrode drift is removed over the period, followed by mean centering. The projection matrix of the actual data is calculated using the PCs generated from the training data.

## III. Results

In this section a brief exposition of our results for validation of different components of the Prototyping Platform for Adaptable Neural Stimulation is presented. The IrOx sensors are first verified to confirm their ability to record both pH and CAPs, recorded through the analogue front-end. Details regarding the analogue front-end are presented in [10]. The results of PCA training and real time pH detection are also discussed here. The closed loop implementation was eventually verified by detecting the change in the rat’s heart rate due to electrical stimulation of vagus from our stimulator, this is shown in [6].

### A. IrOx sensor: pH calibration, CAP recording

Details about the IrOx pH sensor calibrations are given in our previous publication [8]. For CAP recordings we first tested the validity of using an IrOx electrode. We put an IrOx electrode on the same nerve with a Pt electrode for comparison and obrained very good results, see Fig.6.

**Fig. 6.**
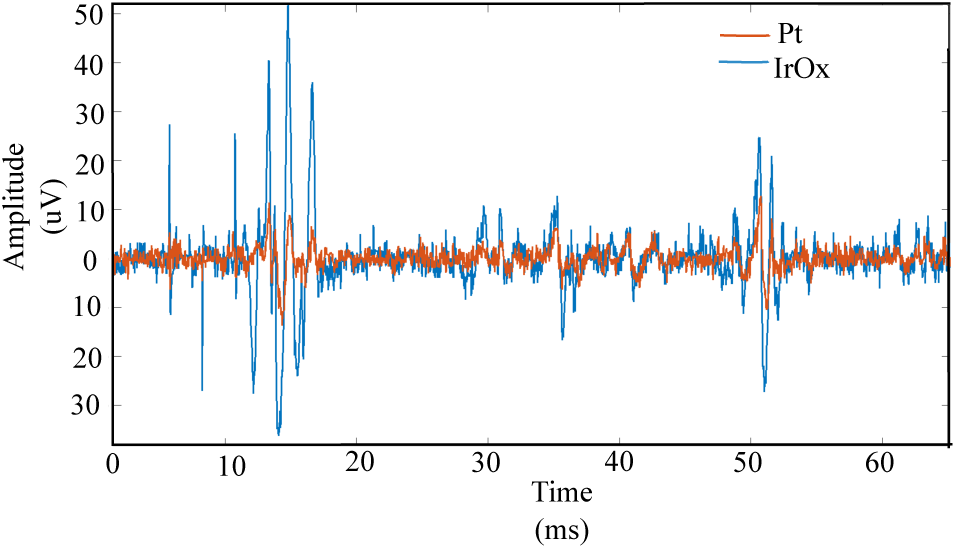
The same CAPs recorded simultaneously by an IrOx and a Pt electrode.

### B. Nerve Classification

The nerve fibre classification can be performed on the basis of CAP conduction velocity. As an example we show some of our experimental results in Fig. 7.

**Fig. 7.**
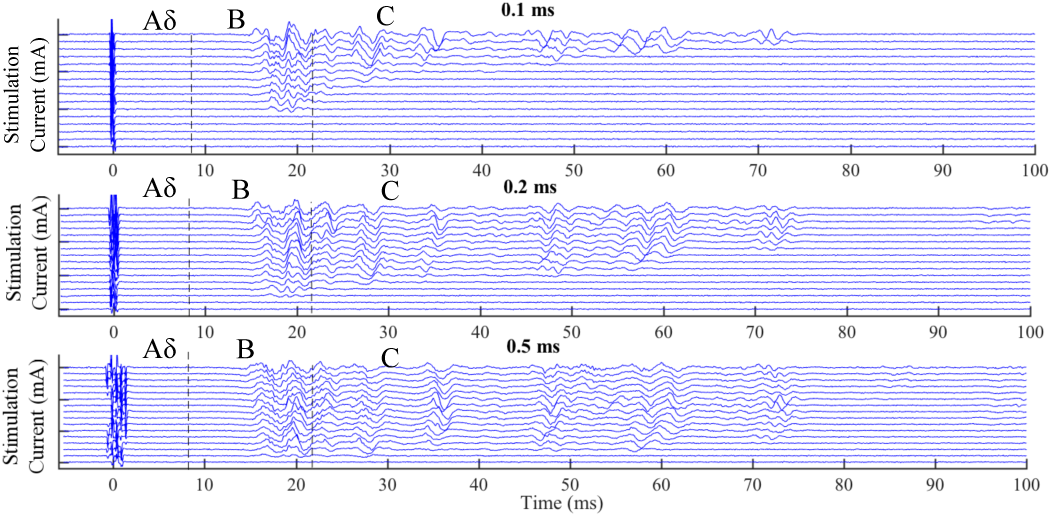
CAPs elicited during *in vivo* stimulation of the cervical part of vagus nerve in a rat and recording in the gastric part. The pulse widths (PWs) shown here are: 0.1 ms (top), 0.2 ms (middle) and 0.5 ms (bottom). More PWs were used in the experiments. There are 16 different current amplitudes for each PW in the span between 0.2 mA-3 mA for PW=0.1 and 0.2 ms, and between 0.1 mA-2 mA for PW=0.5 ms.

### C. Stimulation Strength Determination

The strength-duration curves were used to establish the nerve excitability. Parameters such as rheobase and chronaxie [11] can be calculated from the recordings such as those shown in Fig. 8. Then we can set the stimulation dose (and stimulus profile), depending on which fibre types are targeted.

**Fig. 8.**
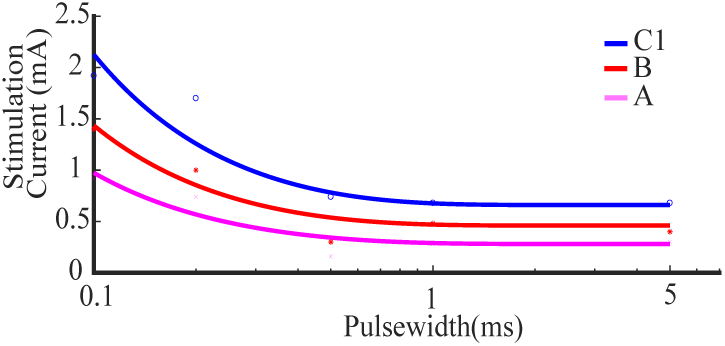
Strength-duration curves for A*δ*, B and C1 fibres.

### D. pH Response Training

The scree plot demonstrates the variance of training matrix data captured by each principle component, shown in Fig. 9. From the scree plot it is apparent that only first three principal components need to be retained, as more than 95% variance in data is accounted for by them.

**Fig. 9.**
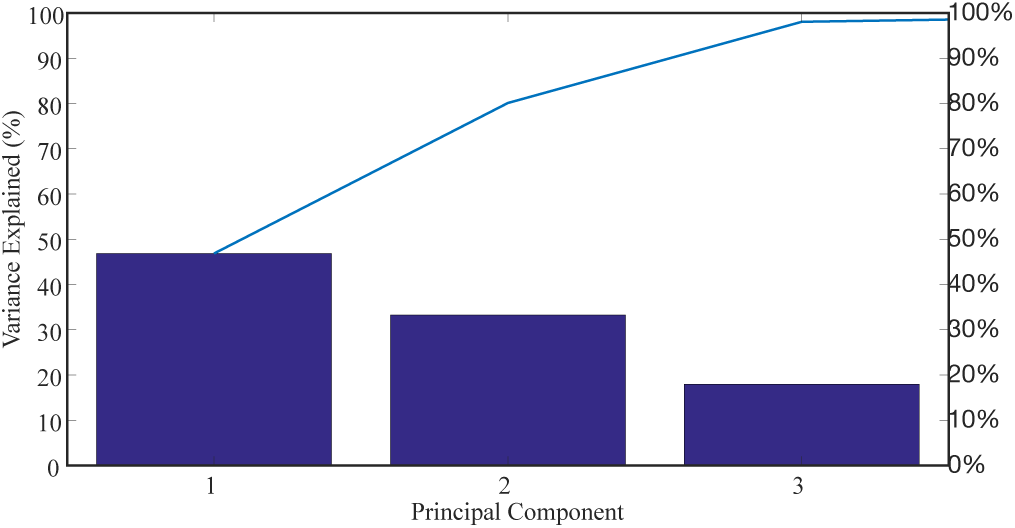
Scree plot of the training matrix shows how much of the variance is covered by the first three principal components: individually (bar plot) and cummulatively (blue line).

### E. Real Time pH Detection

Validation of our algorith is shonw in Fig.10. We show two examples how decision is made on the basis of the residues and *Q*-value, even in the presence of a non-linear background change.

**Fig. 10.**
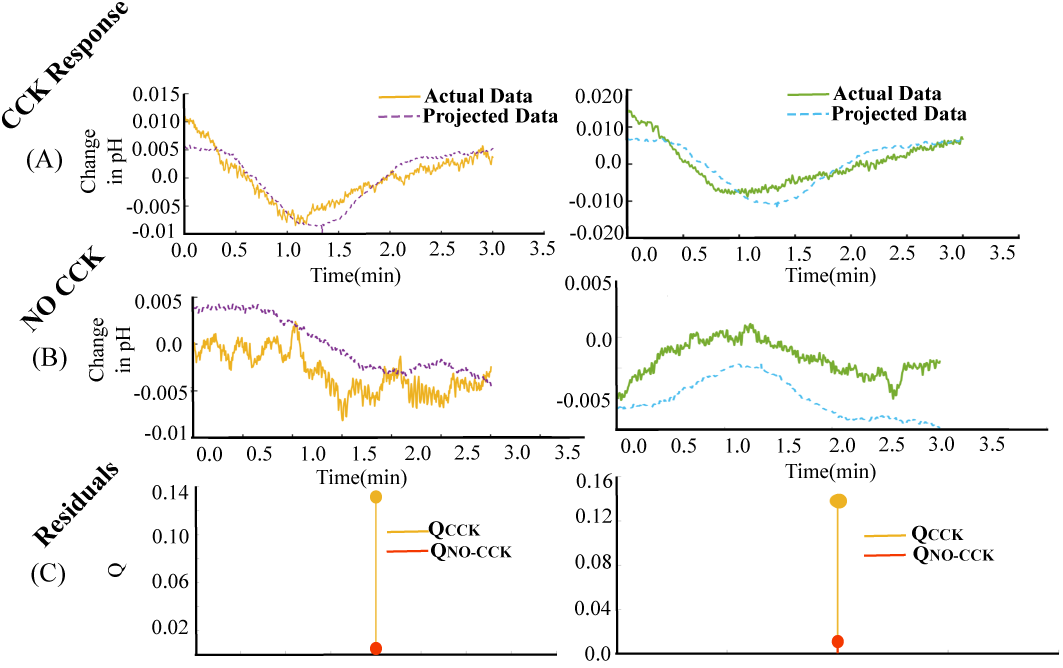
Two examples of the implementaion of our algoritm, shown in left and right columns. (A,B)T he projected data is calculated using the PCs and compared with the actual data. (C) The resiuduals are calculated and *Q*-value shown. *Q <* 0.01 is treated as a response.

## IV. Conclusion

A comprehensive platform for closed-loop neuro-stimulation integrated systems, based on dual chemical and electrical sensors recording, has been created and successfully demonstrated. For the demostration we used a neurometabolic therapy related to vagus nerve stimulation. This platform can be utilised to evaluate closed loop neuromodulation therapies and processing algorithms *in vivo* before translation into an on-chip integrated system for implants.

## Acknowledgment

The authors thank the EU ERC Synergy Grant no. 319818, i2MOVE and UK EPSRC grant EP/N002474/1.

